# Early functional connectivity in the developing sensorimotor network that is independent of sensory experience

**DOI:** 10.1101/2021.06.14.448057

**Authors:** Christine M. Cross, Laura Mediavilla Santos, Nick Whiteley, Karen Luyt, Michael C. Ashby

## Abstract

Neonatal sensory experience shapes development of neural pathways carrying sensory information to the cortex. These pathways link to wider functional networks that coordinate activity of separate cortical regions, but it remains unknown when these broader networks emerge or how their maturation is influenced by sensory experience. By imaging activity across the cortex in neonatal mice, we have found unexpectedly early emergence of coordinated activity within a sensorimotor network that includes whisker-related somatosensory cortex and motor cortex. This network is spontaneously active but is not engaged by sensory stimulation, even though whisker deflection reliably drives cortical activity within barrel cortex. Acute silencing of the sensory periphery ablated spontaneous activity that was restricted to barrel cortex but spared this early sensorimotor network coactivity, suggesting that it is driven from elsewhere. Furthermore, perturbing sensory experience by whisker trimming did not impact emergence or early maturation of spontaneous activity in the sensorimotor network. As such, functional sensorimotor cortical networks develop early, in parallel with development of ascending sensory pathways, and their initial maturation is independent of sensory experience.

## Introduction

Neuronal activity within the developing brain is vital for the correct formation of mature functional neural networks. This activity can be evoked by both external sensory experience and generated spontaneously (Blankenship and Feller, 2010; Katz and Shatz, 1996). Disruption of early neuronal activity patterns often results in the malformation of circuit connections within and between cortical regions (Ackman et al., 2012; Keller and Carlson, 1999; Kirkby et al., 2013; Leighton and Lohmann, 2016), which can then impact maturation of behaviour (Buzsáki, 2010; Harris, 2005; Musall et al., 2019).

Development of functional neuronal networks is a multistage process, beginning with activity independent molecular and genetic guidance, followed by activity dependent processes that can be divided into intrinsically (spontaneous) and extrinsically (evoked) generated events (Blankenship and Feller, 2010; Leighton and Lohmann, 2016; Yamamoto and López-Bendito, 2012). Even early spontaneous activity can be highly organised both spatially and temporally, often propagating through nascent neural connections. Spontaneous activity generated in the periphery is well documented in early stages of sensory network development (Ackman et al., 2012; Babola et al., 2018; Hanganu et al., 2006; Mizuno et al., 2018). For example, spontaneous retinal waves drive activity that propagates all the way to visual cortex and spontaneous twitching of individual whiskers leads to somatotopic activation of somatosensory cortex in neonatal rodents (Ackman et al., 2012; Arroyo and Feller, 2016; Tiriac et al., 2012). These forms of spontaneous peripheral activation of sensory receptors guide initial arrangement of connectivity in preparation for active sensing and the experience-dependent plasticity that it drives. Other sources of spontaneous activity also exist within developing sensory pathways. Spontaneous embryonic thalamic activity that appears before connection to upstream sensory relays shapes patterning of sensory cortical areas (Antón-Bolaños et al., 2019). There are also reports of spontaneous cortical activity that is seemingly independent of the sensory periphery (Siegel et al., 2012; Yang et al., 2009).

The maturation of the circuitry that underpins sensory perception is thought to proceed in a temporal sequence, which follows the order of the synaptic relays that carry the activity from the sensory periphery up to the cortex. As such, more peripheral synapses mature first and are followed, in sequence, by the downstream connections. Indeed, it has been shown that the later development of downstream synaptic connections can depend on the correct, earlier maturation of afferent parts of the pathway. For example, in the sensory pathway carrying whisker information to the primary somatosensory cortex, the brainstem to thalamus synapse matures before the thalamocortical projection into layer 4 of barrel cortex. Recurrent connections within layer 4 then mature in a short window before the maturation of synapses from layer 4 onto layer 2/3 neurons and then layer 2/3 to layer 5, mirroring the sequence of flow of whisker information flow through the mature circuit (Anastasiades and Butt, 2012; Ashby and Isaac, 2011; van der Bourg et al., 2016; Yang et al., 2018). Ultimately, the full maturation of the synaptic pathway depends on plasticity driven by activity from whisker experience at the appropriate time (Erzurumlu and Gaspar, 2012; Fox, 1992).

As with other pathways bringing sensory information from the periphery, whisker-related information integrates into a larger cortical network downstream of the primary sensory area. This connectivity between brain regions underpins the broader functional networks that have been associated with sensory stimulation. In the mature rodent brain, a robust sensorimotor network is characterised by strong functional connectivity between primary somatosensory (S1) and motor (M1) cortex (Aronoff et al., 2010; Chakrabarti and Alloway, 2006; Mao et al., 2011). The integration of information between these regions is vital for sensory perception and motor behaviours (Petersen, 2019). In the mature brain, there is specific functional coactivity of S1 and M1 that occurs both spontaneously (Afrashteh et al., 2020; Mohajerani et al., 2013) and in response to external somatosensory stimuli (Chakrabarti et al., 2008; Ferezou et al., 2007; Manita et al., 2015). However, because of the difficulty of simultaneously recording from multiple regions in the neonatal rodent brain, relatively little is known about when large cortical networks start to be engaged by sensory stimuli or how they mature during neonatal development.

To address these questions, we have used mesoscale calcium imaging to investigate the development of spontaneous and sensory evoked activity across the cortex of neonatal mice. Since these broader cortical networks are driven by sensory experience in the adult brain, we initially hypothesised that their developmental emergence would come after, and would depend on, the maturation of ascending pathways that carry relevant sensory-related activity. However, amongst the large changes in activity patterns that occur through this neonatal period, we found unexpectedly precocious development of a sensorimotor functional network between sensory and motor cortex. This network seems to initially be independent of the classical somatosensory pathway suggesting that its maturation follows an unexpected trajectory.

## Results

### Mesoscale imaging of cortical activity in neonatal mouse pups

To measure the dynamics of neural activity across the neocortex during early postnatal development, we established widefield mesoscale calcium imaging in head-fixed, behaving neonatal transgenic mouse pups expressing GCaMP6. A transgenic strategy was implemented to express the calcium indicator using a cross of Emx1-IRES-cre, which is expressed early in prenatal development, and Ai95D mice, which carry a cre-dependent GCaMP6f gene (Chan et al., 2001; Chen et al., 2013; Kummer et al., 2012). We adopted a breeding strategy using only singly transgenic males that produced pups with GCaMP6 expression restricted predominantly in excitatory neocortical cells (see Methods, Supplementary Figure 1a). To capture the early stages of neocortical functional circuit development, we recorded sensory-evoked and spontaneous activity in pups aged between postnatal day 1 to 9 (P1-P9).

Prior to recording cortical activity, the scalp was removed, and a miniature head-fixation post attached to the skull over the cerebellum under brief (<10 minutes, 1.5-2.5% isoflurane) surgical anaesthesia. The skull was left intact. To avoid any lingering effects of anaesthesia (see Methods, Supplementary Figure 1b-c), analysis of cortical activity was based on data collected at least 60 minutes after removal of anaesthesia. For imaging, pups were head-fixed under a tandem lens fluorescence macroscope in a warmed, nest-like environment (Ratzlaff and Grinvald, 1991). GCaMP6f fluorescence timelapse images of the entire cortical surface were collected at 50 frames per second (Figure 1a). Although the contribution of hemodynamic autofluorescence is small in neonatal mice (Kozberg et al., 2016), we did reliably observe a continual, high frequency (8-10Hz), low amplitude oscillation in fluorescence (Supplementary Figure 1e). This signal, which is consistent with heart rate of neonatal mice, was removed from fluorescence traces using a low-pass 7Hz filter that did not impact detection of lower frequency activity (Supplementary Figure 1d).

**Figure 1.**
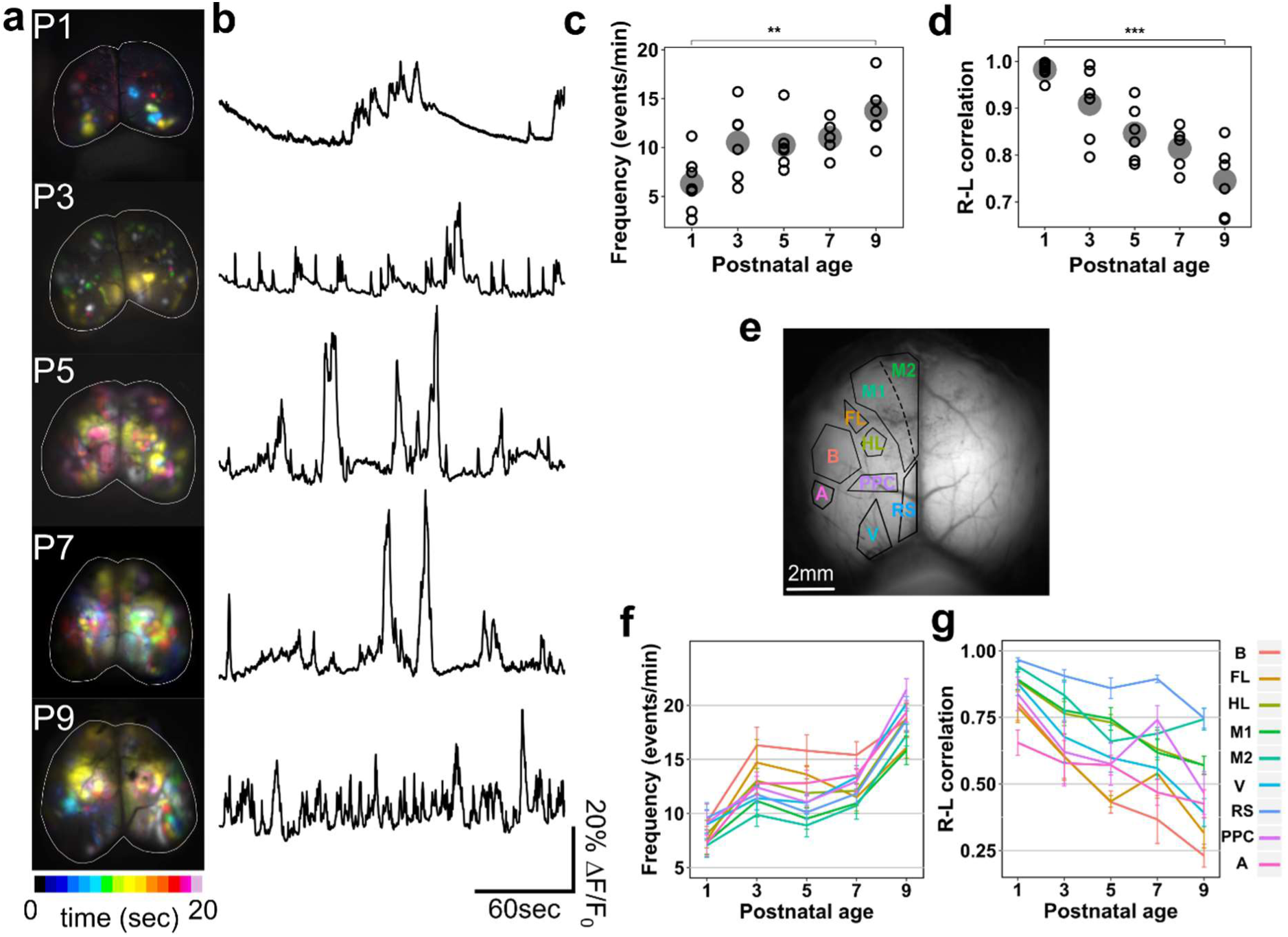
Developmental changes in spontaneous activity across the cortex. a. Time projection maps showing representative spontaneous activity across a 20 second epoch at ages from P1-9. b. Representative fluorescence traces from whole cortex areas (white lines shown on images in (a)). c. The frequency of cortical spontaneous activity increases across postnatal development (p < 0.001, one-way Kruskal-Wallis test). d. Spontaneous activity becomes less bilaterally correlated with postnatal development (p < 0.001, one-way AN OVA). e. Schematic of cortical regions used to parcellate calcium imaging recordings for regional frequencies overlaid on raw fluorescence Image. f. Frequency of spontaneous activity increases with age in all cortical regions with a similar developmental trajectory.

### Spontaneous neonatal cortical activity

Even during periods of behaviourally quiet rest, neuronal networks are still active in both the adult (van den Heuvel and Hulshoff Pol, 2010; Mohajerani et al., 2013; Vanni and Murphy, 2014) and neonatal brain (Ackman et al., 2014; Colonnese and Khazipov, 2012; Doria et al., 2010). Indeed, we observed regionally-localised, ongoing patterns of fluorescence changes across all of the cortical field of view in resting neonatal mice. This activity ranged from highly localised transients or waves to simultaneous, coordinated activation of multiple areas. Time projection maps across 20 seconds of imaging in animals of ages spanning P1-P9 exemplify the spatiotemporal diversity of spontaneous activity (Figure 1a). Spontaneously occurring activity was observed from P1, but the frequency of activity transients across the entire cortex increased with postnatal development (Figure 1a-c) (p < 0.001, one-way Kruskal-Wallis test; n(animals): P1 = 7, P3 = 6, P5 = 6, P7 = 6, P9 = 7). As coordination of neural activity between hemispheres is known to be developmentally regulated, we compared activity in each hemisphere to establish the ability of our approach to identify trajectories of neonatal brain maturation. We found almost complete temporal inter-hemispheric correlation of spontaneous activity just after birth (at P1), and this coordination progressively decreases with postnatal age (Figure 1d) (p < 0.001, one-way ANOVA). At P9 inter-hemispheric correlation was 0.75±0.07, which is still higher than the ∼0.5 seen in the adult cortex (Mohajerani et al., 2010). When broad cortical regions were delineated (based on a scaled brain atlas) and investigated individually (Figure 1e), the trajectory of the age-dependent increase in frequency of spontaneous activity was similar across all regions (Figure 1f). It was evident that there was marked transition in the frequency of spontaneous activity within each individual region, firstly between P1 and P3 and then between P7 to P9 (Figure 1f). In contrast, the broader inter-regional coordination exemplified by inter-hemispheric correlation followed a steadier developmental reduction (Figure 1g). It is notable, however, that the developmental reduction in inter-hemispheric correlation proceeded at quite different rates in different regions, highlighting the variability in maturation of long-range functional networks (Figure 1g).

A variety of spatial motifs of stereotypical spontaneous activity are observed in these recordings, ranging from moments when individual regions were selectively activated to coordinated patterns suggesting broader functional network activity that characterises mature brain function (Chan et al., 2015; White et al., 2011; Wright et al., 2017). To objectively assess the contribution of different activity patterns across developing cortex, we used a non-negative matrix factorization (NMF) approach on aligned images from animals aged P3, P5, P7 and P9 (activity at P1 was so temporally sparse and spatially widespread that NMF was ineffective). NMF yields a representation of the data in terms of spatial activity motifs and their levels of activation over time, which allows identification of common patterns of cortical activation (Figure 2a)(Mackevicius et al., 2019). Furthermore, the number of motifs needed to explain variability in the data allows quantification of the complexity of the underlying coactivation patterns. Running NMF on 3 minute image sequences identified several similar spatial motifs that often recurred in different animals and across recordings (Figure 2b). Several of these common motifs correspond to activity in sensory areas, in particular somatosensory and visual cortex, as might be expected from previous neonatal recordings of neural activity (Figure 2b). There were also more complex, multi-area motifs that were prominent amongst the variety of patterns (Figure 2b). To assess developmental changes in the variety of spatial motifs, we ranked the motifs by the amount they contributed to each recording overall. We then measured the number of motifs required to explain 75% of the total variance of the activity (Supplementary Figure 2). Early in development, fewer motifs accounted for much of the activity, but, with increasing age, the number of identified motifs contributing increased (Figure 2c). This increase in the number of contributing motifs suggests that patterns of spatial activity rapidly become more complex in these neonatal stages.

**Figure 2.**
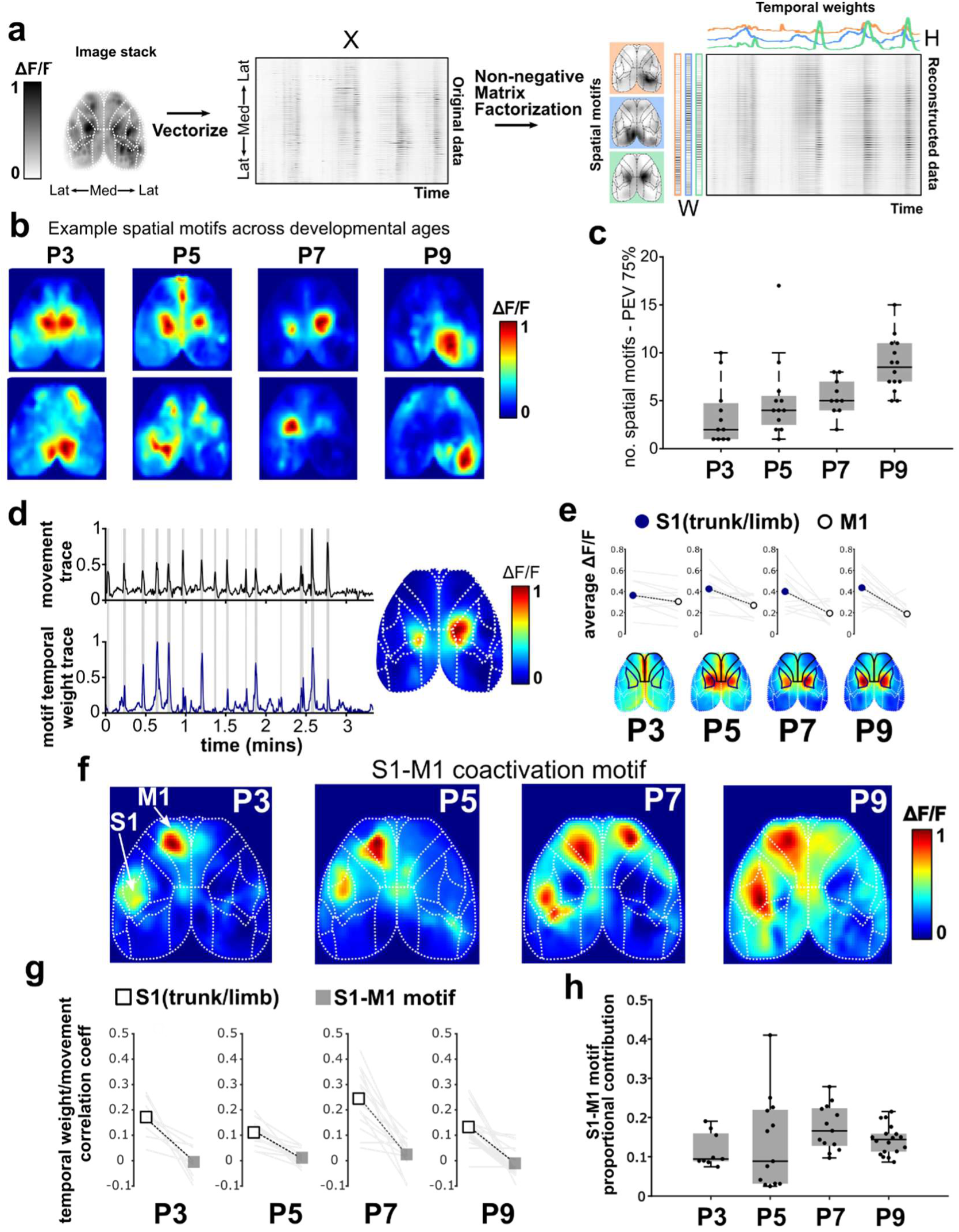
Non-negative Matrix Factorization to identify common spatial motifs of activity across development. a. Process schematic for NMF of image sequences. b. Example spatial motifs of activity extracted by NMF at each postnatal age analysed. c. The average number of inidivudal motifs needed to explain 75% of the variance in the data increases with postnatal age. d. Body movement detected by pressure sensor (upper panel - movement bouts shaded in grey) correlates strongly with the temporal weighting of a particular motif (lower panel) characterized by activity centered on the trunk/limb area of somatosensory cortex (S1) (map overlaid with regional parcellation). e. At all ages tested, the movement-related motif has far greater activity in S1 than in motor cortex (M1). f. Example motifs at each developmental age characterised by simultaneous activity in M1 and in whisker- related S1. g. Relative correlation coefficients between movement sensor and temporal weighting of NMF motifs show that activity in the trunk/limb area of S1, but not with activation of the S1-M1 coactivation motif, is asscoiated with body movement. h. The S1-M1 coactivity motif makes a substantial contribution to overall patterns of activity at each age 180 between P3-P9.

Up to this point, we have grouped all activity regardless of behavioural state. However, all the animals displayed uncoordinated, brief and sporadic movements during the imaging, which were recorded using a pressure sensor placed under the body (Figure 2d, Supplementary Figure 3a). These movements accounted for, on average, ∼10-15% of the total recording duration across all the neonatal ages studied (Supplementary Figure 2b). To assess whether this movement was related to particular patterns of cortical activity, we compared the temporally weighted trace of each extracted NMF motif, which indicates the time and amplitude at which that motif occurs, with the signal from the movement sensor (Figure 2d). In all cases, the strongest correlation with movement was from activity motifs characterised by bilateral activation of a large mid-parietal area that aligns broadly with the location of trunk- and limb-associated somatosensory cortex. However, there was little apparent activity in the more frontal regions where motor cortex is located (Figure 2d). Indeed, across the developmental period we studied, up to P9, movement was much more strongly associated with activity in somatosensory rather than motor cortex (Figure 2e). A disconnect between motor cortex activity and movement in early postnatal rodents has been described previously (Dooley and Blumberg, 2018) and contrasts with the mature brain, in which spontaneous locomotion is associated with activation of large areas of the dorsal cortical surface, including motor cortex (West et al., 2020). However, among the other motifs detected by NMF, there was a commonly occurring pattern at all ages tested between P3 to P9 that did involve the activity in the motor cortex. This motif was characterised by simultaneous activation of the predicted site of primary motor cortex (M1) and primary somatosensory cortex (S1), centred on the predicted site of the barrel cortex (Figure 2e). In contrast to the trunk-limb activity motif described earlier (Figure 2d), this S1-M1 co-activity did not correlate with signal detected by the pressure sensor, suggesting it is not related to movement (Figure 2f). To assess the prevalence of this S1-M1 co-activity pattern, we measured the contribution of this motif to data reconstructed from all the motifs identified by the NMF. The S1-M1 co-activity motif appeared in recordings across all ages and, on average, accounted for 10-20% of the reconstructed data (Figure 2g). This recurring spontaneous coactivation of somatosensory and motor cortex suggests that there may be a sensorimotor functional network even at very early stages of development.

### Sensorimotor network coactivity

To explore the development of activity in specific sensory-related networks more precisely, we investigated the dynamic coordination of whisker-related S1 barrel cortex with other cortical regions. We used seed pixel maps (SPMs) to display cross-correlation between the fluorescence timecourse in S1 barrel cortex and all other pixels across each 3 minute recording. These maps revealed little region-specific correlation at P1. In contrast, in some P3 animals and in all animals at older ages, there was a strong correlation between activity in S1 and an ipsilateral frontal region that aligns with the location of primary motor cortex (M1) (Error! Reference source not found. ai) (Ferezou et al., 2007; Kuroki et al., 2018; Mayrhofer et al., 2019; Vanni and Murphy, 2014). Furthermore, in some of these SPMs, there were also smaller areas of elevated S1 correlation. One of these was just lateral to S1, matching estimated location of secondary somatosensory cortex (S2). The other was just rostral and medial to the M1 area that matches estimated location of secondary motor cortex (M2). Both S2 and M2 are known to have reciprocal connections with S1 in the mature rodent brain (Aronoff et al., 2010; Chakrabarti and Alloway, 2006; Manita et al., 2015). These peaks of elevated correlation suggest that, overall, activity in S1 tends to be temporally coordinated with activity in motor-associated regions. Therefore, even in the early neonatal brain, cortical sensory areas form long-range functional networks reminiscent of those seen in the mature brain (Ferezou et al., 2007).

The consistency of these SPMs increased with postnatal age, with defined regions in S1 and M1 being present in only a few recordings at P3 and becoming progressively more prevalent until all recording blocks produce clear maps at P9. This increasing clarity in S1-M1 correlation suggests there is a developmental strengthening of the sensorimotor functional network.

Reciprocal SPMs centred in M1 revealed similar, but not identical, patterns of correlation (Figure 3aii). The elevated correlation between M1 and S1 was evident in these M1-centered SPMs alongside additional areas of high correlation, such as in contralateral motor regions (Figure 3aii). The fact that S1-centered SPMs and M1-centered SPMs are not the same shows that although spontaneous activity S1 and M1 are coordinated some of the time, they are not always co-active or exclusively coupled.

As SPMs do not reveal information about individual moments in time, we assessed potential coactivity associated with individual spontaneous S1 transients. To do this, we identified peaks in the fluorescence timecourse from S1 and M1 excluding periods associated with movement detected by our pressure sensor. We then assessed whether each peak of S1 activity coincided with a peak in M1 activity (Figure 3b). To capture the possibility of slightly offset timing of peaks in the different regions, a sliding time window was used to identify and measure the proportion of S1 events associated with M1 coactivity at different lags (Figure 3b). Coactivity between S1 and visual cortex (V1) was used as an unrelated cortical comparator. For each recording, we also compared to the proportion of coactivity that could be driven at chance level by measurement of coincidence at a matched number of randomly selected timepoints in both M1 and V1 timeseries. The proportion of coactive events increases as the coincidence window lengthens (Figure 3c). As might be expected, for randomised event timings, this increase manifests as a steadily increasing proportion, with gradient dependent on the frequency of events (Figure 3c). However, at all ages coactivity was higher between S1 and M1 than between S1 and V1 or random timepoints, showing that there is a preferential coordination of individual bouts of activity in S1 and M1 (Figure 3c). There is a rapid increase in the proportion of coactive S1-M1 events up to a ∼100ms duration window, with a gentler gradient of increase after this point in both M1 and V1 that is comparable to random coincidence. This change in gradient in the cortical regions suggests there is an elevated level of biologically driven coordinated events that have their peaks coinciding <100ms apart. Comparing coincident S1-M1 activity within a 100ms window to S1-V1 and random timings in each animal, both M1 and V1 more likely to be co-active with S1 events than predicted by chance (Figure 3d). However, compared to V1, M1 activity is more likely to be coincident with S1 activity at all ages (Figure 3d). The relative proportion of S1-M1 coactivity compared to S1-V1 increases with postnatal age, suggesting that S1 activity is increasingly preferentially associated with M1 co-activation as the brain matures, resulting in a developmental strengthening of this functional network.

**Figure 3.**
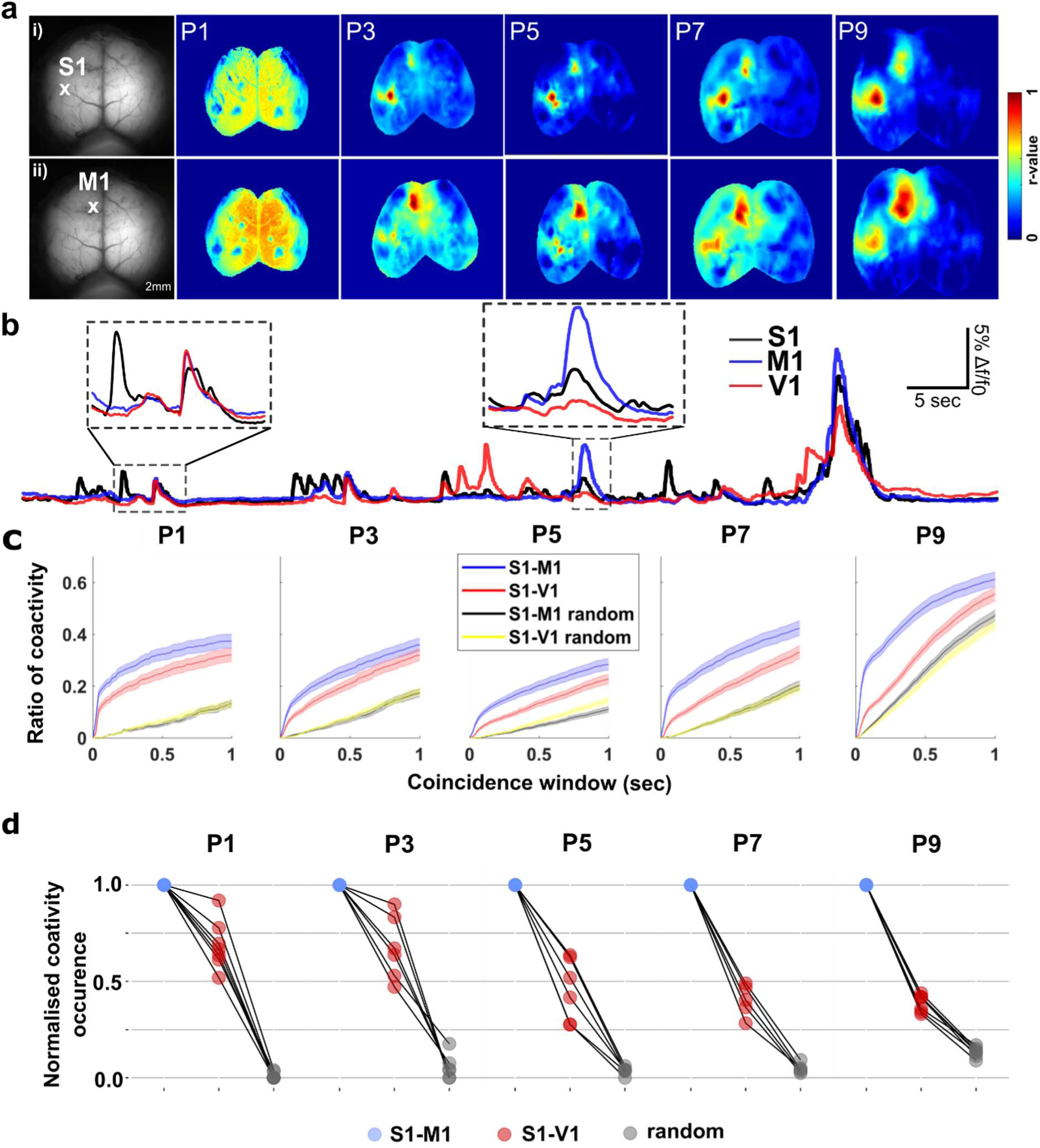
Spontaneous activity in S1 is highly correlated with M1 in the early postnatal brain. a. i) Representative correlation maps from a seedpixel in the barrel cortex from P1-9, during 200s of spontaneous activity, ii) SPM in motor cortex for the same example recordings as i). These show sensorimotor network coordination emerges at P3 and is present across all ages until P9. b. Fluroescence timecourse of pixels located in S1 (black), M1 (blue) and V1 (red). Magnified sections (dashed boxes) show differtnial prevalence and timings of activity peaks in the different regions. c. The proportion of events in S1 that coincide with M1 activity (blue) is higher than V1 activity (red), showing preferential network coordination between S1 and M1. Both regions have higher coincidence than randomly selected time-points, showing that that all spontaneous cortical activity is more coordinated than random. Plot shows mean ±SEM across different coincidence windows. d. Coactivity occurrence within a time window of 100ms of individual animals, normalised to the M1 ratio shows that V1 and random event coactivity is less frequent in all animals.

### Sensorimotor network event categorisation

While we have found evidence of a maturing sensorimotor functional network, many spontaneous S1 events were not accompanied by coactivity in M1 (Figure 4a). To further investigate this, we categorised the spatial organisation of individual S1 spontaneous events. Cortical activation patterns of each spontaneous event in S1 were assigned to being S1-M1 coactive or non-coactive, dependent on whether there was any coincident activity in the M1 region at the peak of the S1 response (Figure 4a).

**Figure 4.**
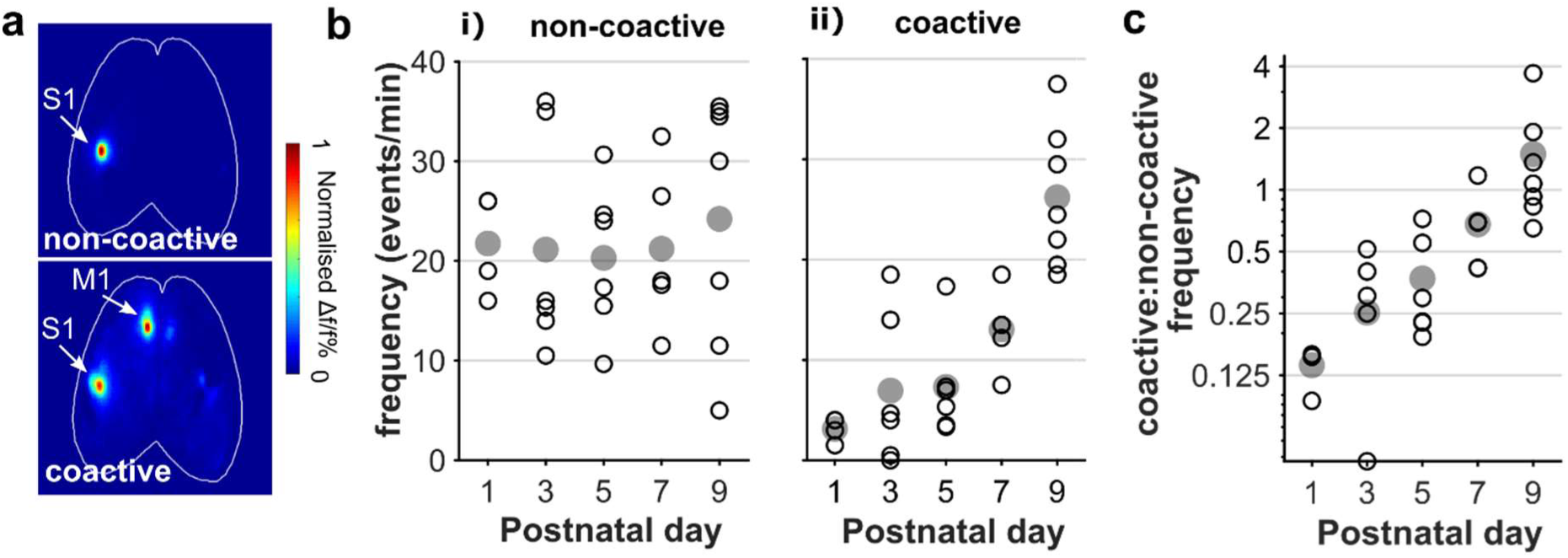
Coordinated S1-M1 spontaneous activity increases across early postnatal development. a. Example activity maps from times coinciding with peaks of spontaneous activity in S1 (barrel cortex) fall into categories characterised by either lack of M1 activity (non-coactive, upper image) or distinct co-activation of M1(coactive, lower image). b. i) The frequency of non-coactive events does not change with development (p = 0.13, one-way ANOVA) but (ii) there is a significant increase in coactive events from P1-9 (p < 0.001 Kruskal-Wallis test). Grey circles are mean of recordings in each animal. c. This differential development in event type results in an increasing ratio of coactive to non-coactive events across early postnatal development (p < 0.001 Kruskal-Wallis test).

The average frequency of non-coactive events did not change between P1 and P9 (Figure 4bi) (p = 0.13, one-way ANOVA, n: P1 = 7, P3 = 6, P5 = 6, P7 = 6, P9 = 7) whereas there was a significant increase in the number of S1-M1 coactive events across development (Figure 4bii) (p < 0.001, one-way Kruskal-Wallis test). This difference in change of frequency means that there is a progressive developmental increase in the ratio of coactive:non-active events between P1 and P9 (Figure 4c; p < 0.001, one-way Kruskal-Wallis test, n: P1 = 7, P3 = 6, P5 = 6, P7 = 6, P9 = 7). These results further support a strengthening on the functional connectivity between S1 and M1 during this early neonatal period.

### Sensory-evoked cortical activity

Whisker deflection during rest stimulates neuronal activity in both S1 and M1 in adult (Ferezou et al., 2007; Mayrhofer et al., 2019; Mohajerani et al., 2013) and juvenile (McVea et al., 2017; Quairiaux et al., 2011) rodents. That type of multi-region activity is reminiscent of spontaneous S1-M1 functional network activity we have found in very early neonatal pups. Therefore, hypothesising that sensory stimulation in neonatal animals would drive activity that engages this S1-M1 network, we deflected whiskers while imaging cortical activity. A single deflection of individual whiskers in neonatal mice aged P3 and P7 activated contralateral barrel cortex (Figure 5a) in a topographically organised manner (Figure 5b), as anticipated from previous studies (Mitrukhina et al., 2014; Yang et al., 2013). However, this S1 activity produced no clear activity in motor regions (Figure 5a) at either age. We assessed the timecourse of fluorescence around whisker deflection in S1 and M1 regions identified as preferentially coactive in spontaneous recordings (Figure 5c). There was a strong S1 activation time-locked to the stimulation but little effect in M1 at either P3 or P7 (Figure 5c; n: P3 = 5, P7 = 7)). This contrasts with the spontaneous coactivity of S1 and M1 in pups of this age (Figures 3 & 4). So why is M1 not activated by whisker deflection when the spontaneous activity suggests that there is already functional connectivity between these two cortical areas?

**Figure 5.**
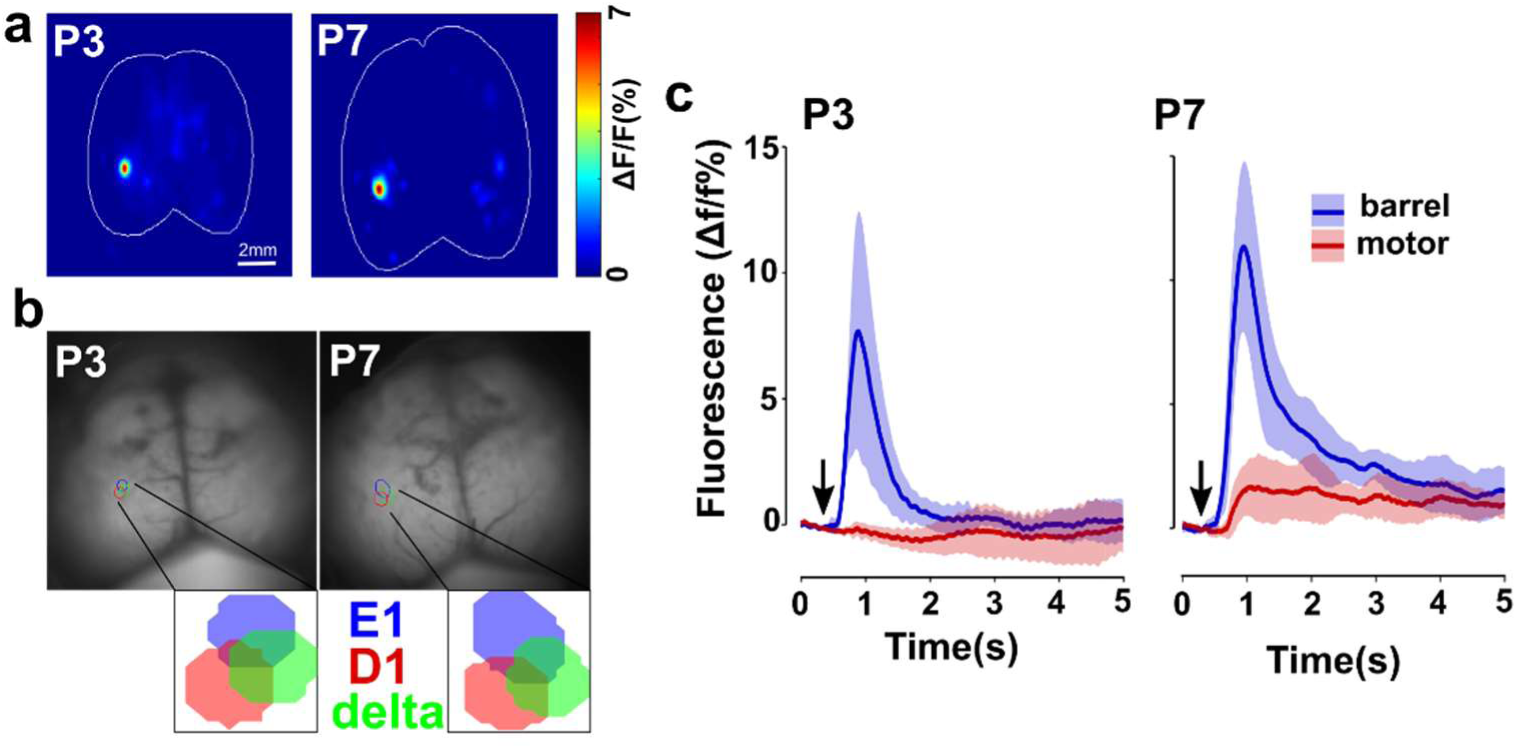
Deflection of a single whisker evokes activation of the contralateral S1 in the first postnatal week. a. Deflection of a single whisker stimulates discrete activity In the contralateral barrel. Example heatmaps of average activity from 20 stimulations from postnatal day 3 & 7. b. This activity is topographically organised. Example regions of activation for deflection of whisker D1 (red), E1 (blue) and delta (green), with closer detail in il). c. Activation of the S1 (blue) is not accompanied by M1 (blue) activation following whisker stimulation (arrow marker). Average (± SD) timeseries of both D1 and E1 (P3, n=5; P7, n=7).

We reasoned that single deflection of a single whisker simply may not be a strong enough driving force to stimulate M1 activity. Therefore, to investigate the consequences of more robust S1 activation, we imaged cortical responses to multi-whisker stimulation in animals aged between P1-9. Deflection of the whiskers again triggered a short-latency activation of contralateral S1 that was already present at P1 (Figure 6a; n(animals): P1 = 7, P3 = 6, P5 = 6, P7 = 6, P9 = 7). The activity was larger amplitude and more spatially widespread than that triggered by single whisker stimulation but was again largely restricted to somatosensory cortex (Figure 6a; single:multi-whisker % of cortex activated - P3, 0.30:0.85; P7, 0.48:1.26). In particular, this sensory stimulation did not result in robust activation of M1 at any age between P1 and P9 (Figure 6b).

**Figure 6.**
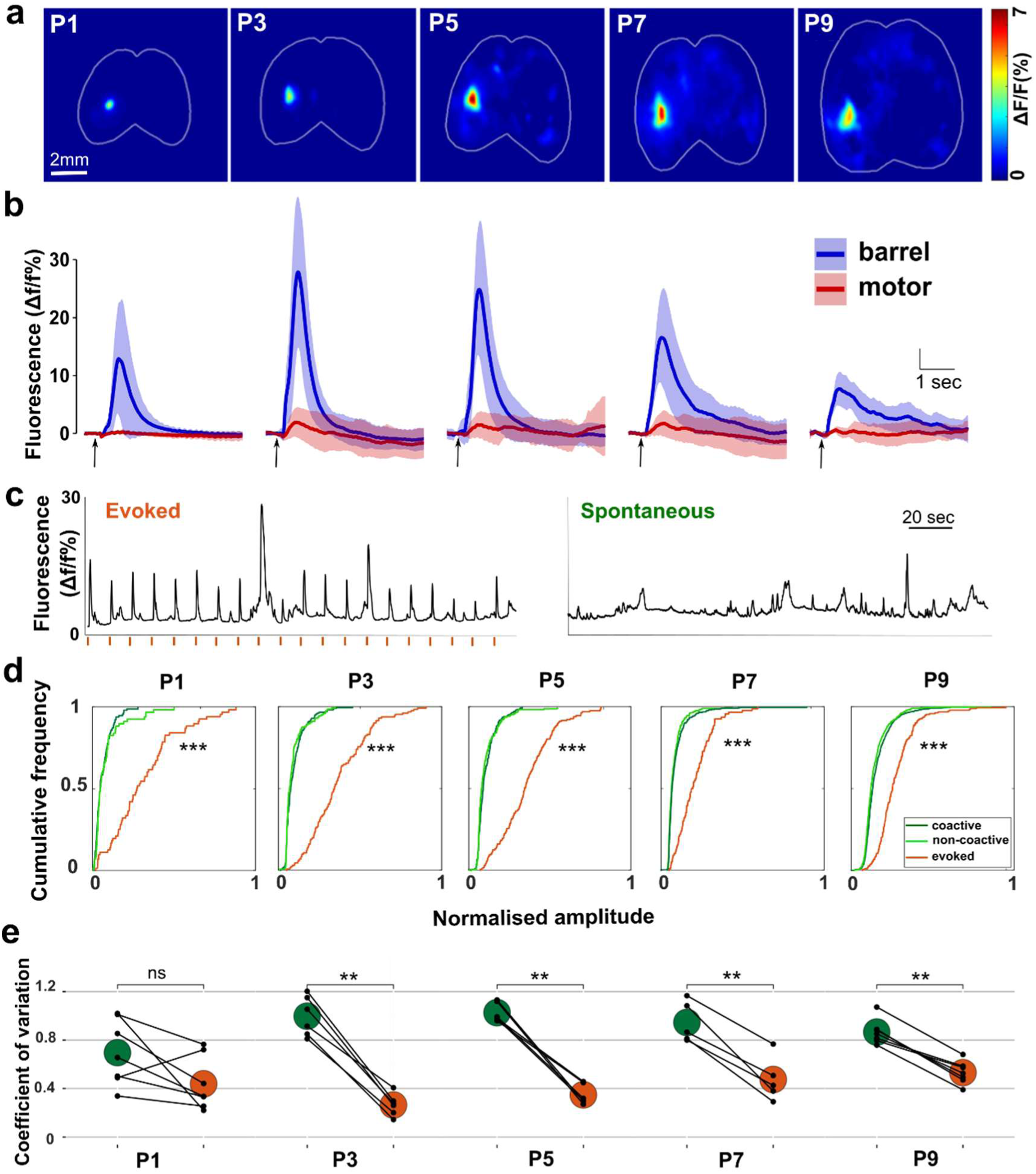
Deflection of multiple whiskers evokes activity in contralateral S1 but not M1 in early postnatal mice. a. A single deflection of multiple whiskers simultaneously results in activation of the contralateral barrel cortex (S1) from P1 to P9. Example heatmaps of average activity from 20 stimulations. b. Activation of S1 (blue) following whisker stimulation (black arrow marker) is not accompanied by M1 (red) activity. Average (± SD) timeseries. c. Example events of both whisker evoked (orange lines mark stimulation times) and spontaneous activity in S1. d. Amplitudes of spontaneous S1 activity (both coactive and non-coactive) is smaller than whisker evoked events from P1-P9 (p < 0.0001, K-S test). Plot showing the cumulative frequency of amplitude of individual spontaneous (green) and evoked (orange) events in S1, normalised to maximum amplitude. e. The amplitudes are more variable for spontaneous events, with the coefficient of variance in spontaneous recordings being significantly higher than stimulated events from P3-9 (p < 0.01, two-way repeated measures ANOVA- paired within animal), (n: P1 = 7, P3 = 6, P5 = 6, P7 = 6, P9 = 7).

We directly compared the properties of sensory-evoked and spontaneous events in S1 for each animal. Stereotyped responses were reliably evoked by each multi-whisker stimulus (Figure 6c). By comparison, spontaneous activity within the same region was much more variable in amplitude and kinetics (Figure 6c). Indeed, comparison of their relative amplitude showed that most spontaneous events were, on average, smaller than evoked events across all ages (Figure 6d). This was the case for both S1-M1 coactive and non-coactive events, which had similar amplitude distributions (Figure 6d). Notably, though, some spontaneous events were of similar amplitude to evoked responses. Variability (quantified as coefficient of variation) of spontaneous event amplitude was higher than evoked responses in all but one animal. Comparison across development showed that there was significantly more variability in spontaneous activity than evoked activity from P3 up to P9 (p< 0.01) (Figure 6e).

The spatiotemporal variability in the S1 spontaneous event and the differential spatial properties of whisker-evoked suggests that there may be distinct drivers of spontaneous activity within the sensorimotor network. We have shown that passive whisker simulation can drive S1 activity, but it is unclear whether S1-M1 co-activity also relies on the classical activation of peripheral sensory pathways.

### Silencing the sensory periphery alters spontaneous cortical activity

To assess the contribution of peripherally generated spontaneous activity to the sensorimotor cortical network, we acutely silenced the whisker-related sensory drive. To achieve this, we measured evoked and spontaneous cortical activity in P7 animals before and after the local anaesthetic lidocaine was injected into the right whisker pad.

Lidocaine injections successfully silenced the right whisker pad, consistently producing almost complete inhibition of the contralateral S1 activation driven by whisker stimulation (Figure 7abc). In contrast to whisker deflection-evoked activity, a substantial amount of spontaneous activity was still apparent even after silencing of the whisker pad. The lidocaine injection did reduce the frequency of spontaneous events in contralateral S1 (p > 0.001, paired t-test, n = 7) and in M1 (p = 0.019, paired t-test, n = 7), but ∼60% of S1 and ∼85% of M1 activity remained (Figure 7d). As expected, there was no change in the frequency of spontaneous activity in the ipsilateral hemisphere (S1 – p = 0.061; M1 – p = 0.383, paired t-test, n = 7) (Figure 7d). Therefore, it appears that activity from the periphery drives some, but not all, of the activity in the sensorimotor cortical network and its silencing has the greatest impact on activity in S1. To assess whether peripheral drive is associated with particular modes of spontaneous cortical network activation, we measured the effect of lidocaine injection on the prevalence of S1 non-coactive and S1-M1 coactive events in the contralateral hemisphere. Silencing of the whisker pad caused almost complete inhibition of S1 non-coactive events (p < 0.001, paired t-test, n=7) but only a partial (∼50%) reduction in the frequency of S1-M1 co-activity (p = 0.006, paired t-test, n=7) (Figure 7e). This resulted in a shift towards coordinated S1-M1 activity, dramatically increasing the ratio of coactive to non-coactive events in the contralateral hemisphere (Figure 7f) (p = 0.012, paired t-test). There was no change in the frequency of either coactive or non-coactive events in the ipsilateral hemisphere (non-coactive: p = 0.345; coactive: p =0.206, paired t-test, n=7).

**Figure 7.**
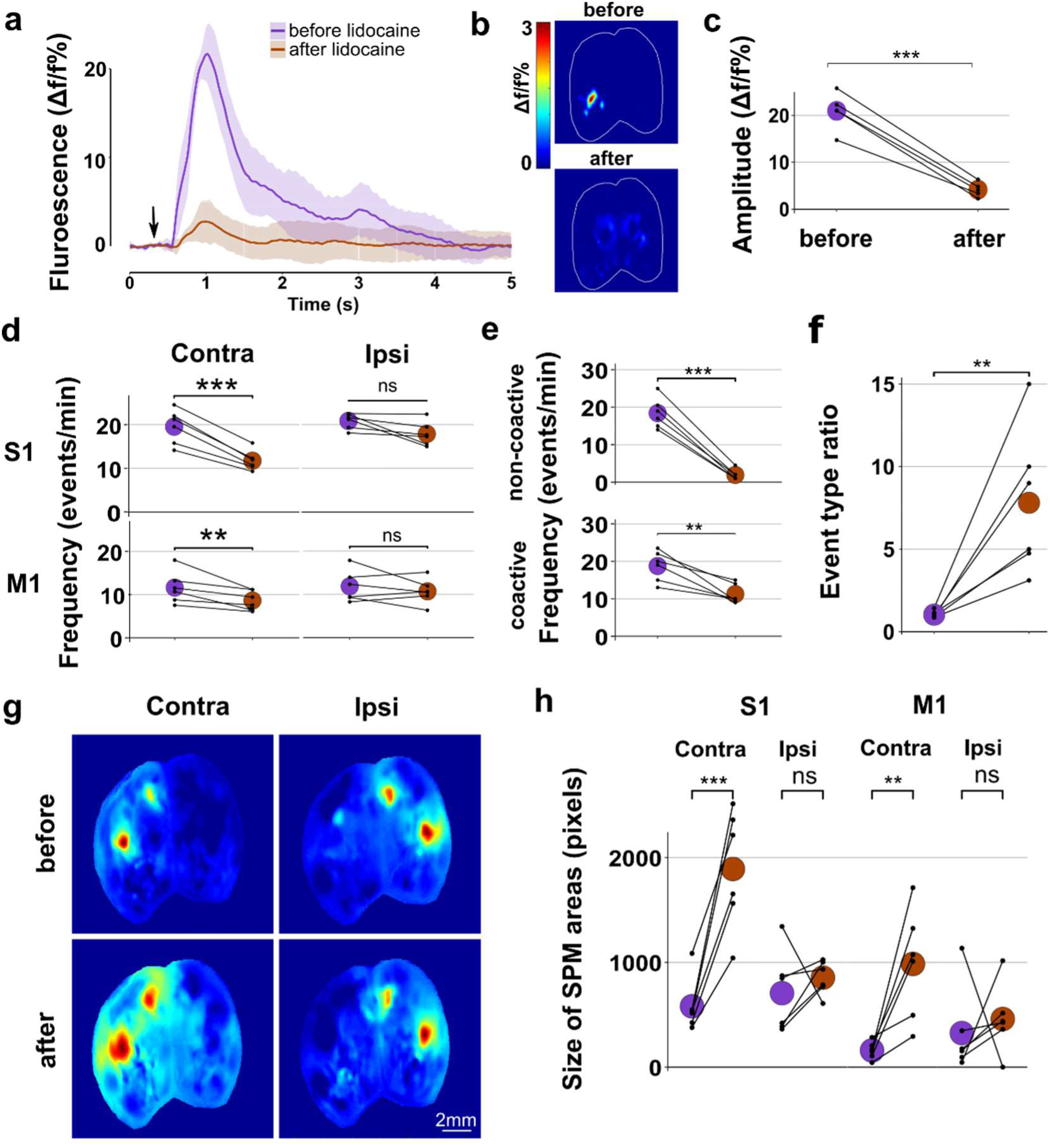
Spontaneous cortical activity is partially driven by the periphery. a. Silencing of the whisker pad with lidocaine injection eliminated cortical activation by whisker stimulation. Plot of average (+SD) activity of S1 in the 5s period following stimulation (black arrow marker). b. Heatmaps showing the loss of specific spatial activation. c. The peak amplitude of the response in S1 is significantly reduced following lidocaine (p < 0.001, paired t- test) d. There is a reduction in spontaneous activity in left (contralateral to injection) S1 after lidocaine administration (p < 0.001, paired t-test - grey circles = mean) and in left M1 (p = 0.019) but not in the right S1 (p = 0.0611) or in M1 (p = 0.383). e. The spatial patterns of individual events are altered by lidocaine administration. In the left (contralateral) hemisphere there is a significant decrease in both S1 only (p < 0.001) and S1-M1 coactive (p = 0.006) events. f. This differential change in event frequency between types results in a significant increase in the ratio of coactive to non-coactive events (p = 0.012, paired t-test) g. An example seedpixel maps centred on S1 showing an increase area of correlation after lidocaine in contralateral hemisphere, h. There is a significant increase in the size of correlated area in both contralateral S1 (p<0.001, paired t-test) and M1 (P < 0.01) but not in the ipsilateral hemisphere (S1 - p = 0.478; M1 - p = 0.655).

This preferential effect of lidocaine treatment on S1 localised activity also changed the overall spatiotemporal properties of correlations in spontaneous cortical activity, which was evident in S1-based SPMs (Figure 7g). Silencing the whisker pad increased the average correlation between contralateral S1 and M1, without affecting the ipsilateral hemisphere (Figure 7g). Also, there was an increase in the size of highly correlated areas in both contralateral S1 (p < 0.001, paired t-test) and M1 (p < 0.001), but not in the ipsilateral hemisphere (S1 – p = 0.478; M1 – p = 0.655) (Figure 7h). Overall, these effects of peripheral silencing suggest that spontaneous activity that is restricted to S1 comes largely through the classical sensory pathway. However, broader sensorimotor network activations involving coordination of S1 and M1 are less likely to arise from peripheral drive. Indeed, the majority of the spontaneous activation of the sensorimotor network is driven largely independent of the sensory periphery in these developing animals.

### Sensory experience dependence of functional network development

It is well-established that early life sensory experience is necessary for the accurate formation for many sensory-driven neuronal networks (Colonnese & Khazipov, 2010; Daw et al., 1992; Feldman, 2009; Kevin Fox & Wong, 2005; Hensch, 2004a; Weller & Johnson, 1975). Given the early appearance of coordinated S1-M1 activity (Figure 3) and with it being different from sensory-evoked activity (Figures 5 & 7), we investigated whether spontaneous activation of the sensorimotor network is also influenced by neonatal sensory experience. We unilaterally trimmed all the whiskers each day from birth because developmental perturbation of whisker experience is known to alter the maturation of the neural pathways carrying sensory information to and within S1 (Ashby & Isaac, 2011; Feldman & Brecht, 2005; Fox, 1992). We investigated the effects of perturbing sensory experience by comparing cortical activity in animals that had undergone unilateral daily whisker trimming from birth to sham-trimmed littermates, imaging P3 and P7 animals (Figure 8a).

**Figure 8.**
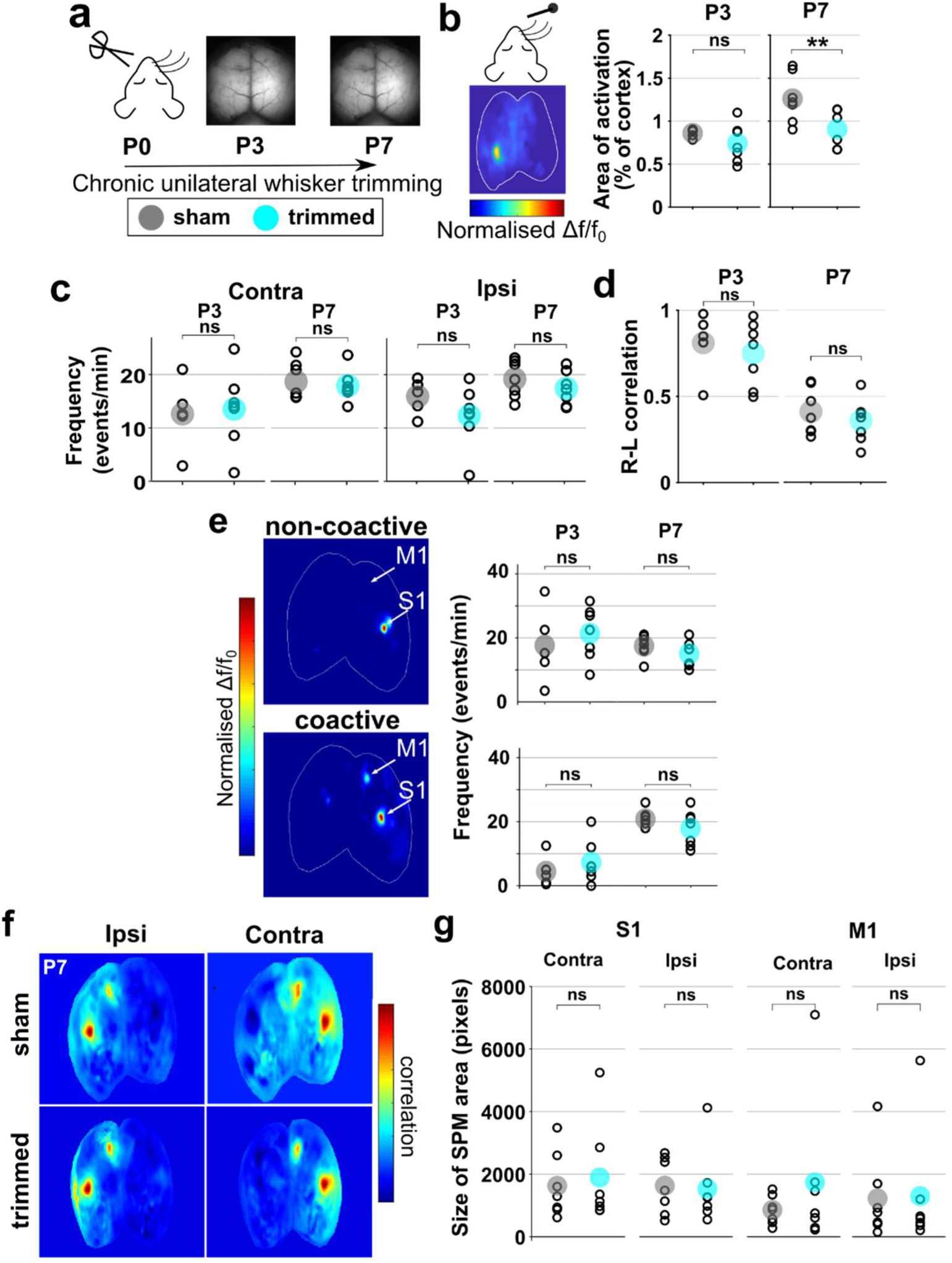
Chronic sensory deprivation during early postnatal development does not alter the temporal or spatial properties of spontaneous cortical activity in the sensorimotor cortex. a. Experimental timeline showing daily unilateral whisker trimming from PO to P7, with imaging of spontaneous and spared whisker-evoked activity in P3 and P7 animals. b. Deflection of spared whiskers evokes activity in barrel cortex ipsilateral to trimmed whiskers, as shown in representative image. The area of cortex activated is smaller in whisker trimmed animals than sham controls at P7 (p = 0.03), but not at P3 (p = 0.27). c. The frequency of spontaneous events in S1 is not altered following chronic sensory deprivation in the contralateral (p = 0.99, two-way ANOVA, n = 7/group) or ipsilateral (p = 0.116, two-way ANOVA) hemisphere, at P3 or P7. d. The correlation of activity between the right and left barrel cortex is not altered in whisker trimmed animals compared to controls at either age (p = 0.327). e. The frequency of individual spontaneous S1 non-coactive and S1-M1 coactive events in the contralateral cortex is not altered by whisker timming (non-coactive; p = 0.899, coactive; p = 0.991, two-way ANOVA). f. Example seedpixel maps centered on S1 contralateral and ipsilateral to whisker trimming from P7 animals. g. At P7 there is no significant difference in the size of correlation area for S1 (contra- p = 0.946; ipsi - p = 0.818, Wilcox test) or M1 (contra- p = 0.597; ipsi - p = 0.946, Wilcox test)

As we could not confirm the effect of trimming on the response to whisker deflection (because the whiskers were still trimmed at the time of recording), we took advantage of the fact that chronic unilateral whisker trimming has been shown to affect spatial properties of responses to deflection of the untrimmed whiskers on the other side of the face (Glazewski et al., 2007). Therefore, we measured the response to whisker deflection of the spared whiskers in groups of animals at P3 and P7. Multi-whisker deflection triggered reliable activation of contralateral S1 (NB – ipsilateral to the trimmed whiskers) similar to that described in earlier (Figure 6). However, by P7, the size of the activated area was significantly smaller in trimmed animals than in controls (Figure 8b; P3 – p = 0.27; P7 – p = 0.03, T-test). This shows that neonatal whisker trimming effectively altered the development of sensory-evoked responses, as expected.

Next, we compared spontaneous activity in the sensorimotor network in the trimmed and sham animals. The frequency of spontaneous events in contralateral or ipsilateral S1 was not altered by perturbation of sensory experience at either age (Figure 8c) (contra - p = 0.99, ipsi – p = 0.116, two-way ANOVA, n = 7/group). Correlation of spontaneous activity between the hemispheres was also unaffected by the whisker trimming (Figure 8d) (p = 0.327), similar to the developmental decorrelation observed in the previous dataset (Figure 1d&g). When the spatial organisation of these S1 spontaneous events was categorised, we found that whisker trimming had not changed the frequency of non-coactive or coactive events in either the contralateral (S1 – p = 0.899, S1-M1 – p = 0.991) or ipsilateral (S1 – p = 0.636, S1- M1 – p = 0.182) hemisphere (Figure 8e). The preferential coactivity in S1 and M1 was also similarly clear in S1 SPMs from P7 sham and whisker-trimmed animals (Figure 8f). To check whether there were more subtle changes in cross-correlation within the SPMs, we measured the size of highly correlated regions in S1 and M1. There was no change in the area of the correlated S1 and M1 regions in either the contralateral (S1 – P = 0.946; M1 – p = 0.597, Wilcox test) or ipsilateral (S1 – P = 0.818; M1 – p = 0.946) hemisphere following chronic whisker trimming (Figure 8g). Overall, these data suggest, in contrast to sensory-evoked activity, that neonatal development of spontaneously generated activity in the sensorimotor network proceeds unabated even in the absence of normal whisker experience.

## Discussion

Using widefield calcium imaging of the brain in mice pups (P1-P9), we have investigated the neonatal development of both spontaneous and sensory stimulus-evoked cortical activity. We found that functional connectivity between regions of the sensorimotor cortex is present from the beginning of the first postnatal week. This network coordination of whisker-related somatosensory and motor cortex is present during spontaneous activity but is not evoked by sensory stimulation at these young ages, even though sensory-driven activity does reach the cortex. This suggests there is a precocious network linking cortical regions that is, at least initially, largely independent of the maturation of the classical sensory pathway. Indeed, acute silencing of the whisker pad had relatively little effect on these spontaneous network activations compared to whisker-evoked or spontaneous activity that is restricted to barrel cortex. Furthermore, by trimming whiskers from birth, we have shown that the formation and initial maturation of this spontaneous sensorimotor network activity is independent of neonatal sensory experience.

### Movement related activity

At the neonatal ages we have studied, movement of the body and limbs is poorly controlled, reflexive or spontaneous (i.e. twitches). We did not find much activity in motor cortical areas that was associated with periods of movement. This aligns with previous findings, which suggest that M1 control of movement starts later in development (Chakrabarty and Martin, 2000; Dooley and Blumberg, 2018; Young et al., 2012). Indeed, in our recordings, the vast majority of movement-related activity was found in centred in the trunk and limb-associated somatosensory cortex, suggesting that it is driven by the sensory consequences of the body motion (Figures 2 & S2)(Khazipov et al., 2004).

It has been shown previously that electrical somatosensory stimulation or myoclonic twitches during sleep can elicit responses in neonatal motor cortex neurons (An et al., 2014; Tiriac et al., 2014). It is possible that this type of reafferent signal is present in the motor areas in our recordings, as there are some hints of activity there (Figure S2), but they are dwarfed by the barrage of somatosensory activity. It is unlikely that our pressure-based sensor detected all forms of movement, particularly localised events such as myoclonic twitches (unless they result in touch of the body onto the sensor). Therefore, cortical activity associated with isolated twitches will be part of what we have termed “spontaneous” activity in our analysis. Whilst we have focussed in this study on periods that are not associated with gross body movement, the cortical activity associated with movement is likely to play a significant role in the development of cortical circuitry.

### Development of spontaneous activity

Spontaneously generated activity is present across the cortex from P1, and the frequency increases with developmental age (Figure 1). Indeed, the relatively low frequency, intermittent activity seen at younger ages is reminiscent of the discontinuous EEG traces that characterise prenatal human babies (Tolonen et al., 2007; Vanhatalo and Kaila, 2006). By P9, there is almost continuous cortical activity, ranging from brief, spatially discrete transients to large travelling waves (Figure 1). Again mirroring EEG activity in human preterm babies, activity in the youngest animals was highly regionally-correlated across hemispheres (Figure 1). Despite the increasing frequency of activity, this inter-hemispheric regional coordination declined with age, perhaps reflecting the development of lateralized specialisation.

Whisker stimulation evokes activity in S1 from P1 (Figure 6a) and we found topographical organisation of individual barrels at P3 (Figure 5b). This agrees with previous findings of organised evoked activity from birth in rodents (Yang et al., 2013). This early S1 whisker-stimulated activity is not accompanied by consistent activation of M1, or any other areas, as is found in the more mature brain (Ferezou et al., 2007). As an isolated result this might suggest that functional connectivity between S1 and M1 is not yet established in this early neonatal period. However, when spontaneously occurring activity was analysed, preferential coactivity between S1 and M1 is clearly present from as early as P1 (Figures 2&3) (McVea et al., 2017). Furthermore, in the time-averaged activity SPMs from many animals, spontaneous S1 activation is highly correlated with distinct areas that align with projected location of M1, M2 and S2 (Figure 3). This suggests that a functional sensorimotor network between these areas is already established very early in development. The strength of correlation between spontaneous activity in these sensorimotor areas increases during the first postnatal week, indicating that the network does undergo a postnatal maturation process. Nonetheless, even by P9, even though deflection of the whiskers reliably drives activity in S1, it fails to engage this broader sensorimotor network. This suggests that the sensory pathway is somehow separated from this sensorimotor network and that they are developing in parallel. In line with the idea that there are parallel networks in early S1, when we analysed cortex-wide activity associated with individual spontaneous events there, we found they fell into distinct types of spatial motifs. While there was spontaneous activation of S1 in patterns similar to those evoked by whisker deflection with activity largely restricted to S1 barrel cortex, there were also many spontaneous events where S1 and M1 were active simultaneously (Figure 4a). The S1-M1 coactive events became relatively more prevalent with age (Figure 4bc), perhaps reflecting a strengthening of the connections between those sensorimotor areas or an increased likelihood of triggering activity within that network.

### What are the drivers of spontaneous activity?

The varying intra-regional functional connectivity during spontaneous and evoked activity raises the question of whether there are different upstream drivers of these activity patterns. We know that whisker deflection drives activity from mystacial sensory receptors to the cortex via brainstem and thalamus (the classical sensory pathway) (Petersen, 2019). This evoked activity is blocked by local anaesthesia of the whisker pad (Figure 7).

Spontaneous neuronal activity originating in the periphery is well documented in the developing sensory networks, including visual, auditory, and somatosensory (Ackman et al., 2012; Akhmetshina et al., 2016; Babola et al., 2018; Mizuno et al., 2018; Torborg & Feller, 2005; Wang et al., 2015). However, it is not the only source of activity. In the developing visual cortex, as well as waves of activity originating in the retina, there are also cortically generated events (Siegel et al., 2012) and silencing of the somatosensory periphery does not eliminate all S1 activity (Yang et al., 2009). When we silenced the whisker pad there was a ∼40% reduction in spontaneous activity in the contralateral S1 (Figure 7d). These events may be triggered by myoclonic whisker twitches or spontaneous activation of sensory receptors (Gómez et al., 2021; Tiriac et al., 2012). The remaining 60% of spontaneous activity in S1 has an origin outside of the peripheral sensory neurons. The S1 activity left is preferentially coactive with M1 and this distinction may tell us something about the origins of the remaining spontaneous activity. In the mature brain there is substantial direct intracortical connection between S1 and M1 (Aronoff et al., 2010; Chakrabarti et al., 2008; Ferezou et al., 2007; Mao et al., 2011). These connections are between S1 layer II/III (Hooks et al., 2011) and M1 layer V (Mao et al., 2011). Layer IV of S1 rapidly develops around the end of first postnatal week (Arakawa and Erzurumlu, 2015; Ashby and Isaac, 2011; López-Bendito and Molnár, 2003) with layer II/III following behind maturing around P12 (Stern et al., 2001). As well as immature inter-regional connectivity in the first postnatal week there is a lack of direct glutamatergic intracortical S1 to M1 connection at P6 (McVea et al., 2017). However, M1 to S1 connectivity is present during this early postnatal period, in both excitatory and inhibitory neurons (Vagnoni et al., 2020). Coactivity between S1 and M1 in the developing cortex is known to be bidirectional, and it has previously been found that peripheral silencing of the paw preferentially eliminates S1 to M1 spontaneous activity (An et al., 2014). The timescale of the calcium indicator used for this study means we do not have the temporal resolution to know for sure whether coactive events originate in S1 or M1, but the source being outside of S1 could explain the preferential resilience of coactive events to peripheral sensory neuron silencing.

Another possible source of spontaneous activity is the thalamus. The first 9 postnatal days are rapidly developing stage of thalamocortical connectivity in S1. The first postnatal week features TC migration and a critical window of plasticity in S1 layer IV (Crair and Malenka, 1995; Lu et al., 2001). Patchwork barrel-related spontaneous activity is evident in layer IV neurons before P9 and this is almost entirely driven from the sensory periphery via thalamocortical relay (Mizuno et al., 2018). This aligns with the largely S1 non-coactive events we have described, which also were entirely dependent on drive from the whisker pad (Figure 7). However, we also found spontaneous activity, largely S1-M1 co-activity, that did not come from the whisker pad (Figure 7). Overall this suggests that the spontaneous co-activity in the larger sensorimotor network at these young ages may not involve layer IV. The widefield imaging technique we used for this study captures neuronal activity from across the upper cortical layers including layer IV. As such, it is tempting to suggest that the different types of spontaneous activity patterns we have described may be segregated in different cortical layers. Although layer IV is the major TC input layer in S1, there are direct thalamic connections and long-range intra-cortical connections to layer II/III that could serve as independent drivers of spontaneous activity (Viaene et al., 2011). As the local connections between different layers mature, activity patterns may become less segregated. Indeed, after P9, there is a major functional switch, with spontaneous activity in barrel cortex layer IV becoming independent of the subcortical sensory pathway (Nakazawa et al., 2020). Whisker-driven sensory information reaches barrel cortex via both first order (ventroposteromedial thalamic nucleus (VPM)) and higher order (posterior thalamic nucleus (POM)) thalamic regions (Petersen, 2019). In adult rodents, it is known that POM (but not VPM) also has a direct synaptic connection to M1 (Chakrabarti and Alloway, 2006), raising the possibility that spontaneous POM activity could simultaneously drive S1 and M1. However, the neonatal development of these pathways is not yet well described.

As well as targeting multiple cortical layers, TC connections that are developing during this early postnatal period have been found to project to multiple cortical regions, some of which are developmentally transient connections and some which persist throughout adulthood (Henschke et al., 2015, 2017, 2018). These cross-modal connections may be responsible for coordinated cortical activity such as between S1 and M1 found in our study. It is known that these cross-modal connections have an important functional role in the development of individual sensory regions, with loss of one sense resulting in remodelling of TC connections in another (Dooley and Krubitzer, 2019; Moreno-Juan et al., 2017). During early postnatal development the subplate is an important relay area for peripheral to cortical connection, and like the thalamus, the subplate has connections spanning across the cortex (Hoerder-Suabedissen and Molnár, 2012, 2015). It is known that subplate neurons generate spontaneous activity which mediates activity in the developing thalamus and cortex (Hanganu et al., 2001) and is vital for correct development of S1 (Tolner et al., 2012).

The amplitude of spontaneous activity was more variable than in evoked S1 activations (Figure 6e). This could indicate that what is being observed is a heterogeneous population of spontaneous events that come from varied sources. This contrasts with sensory-evoked responses, which originate from a single source of consistent drive and so result in lower variability. While there are S1 only spontaneous events that are originate in the periphery (Figure 7), the majority of S1-M1 coactive events are driven internally from one or more of the above sources.

### Experience does not shape the early development of spontaneous functional sensorimotor connectivity

Given the sensitivity of ascending sensory pathways to experience in the early stages of life (Erzurumlu, 2010; Hensch, 2004), we might anticipate that development of downstream intra-cortical functional networks might also be affected by disruption of sensory input. However, chronic unilateral whisker trimming in the first postnatal week did not alter frequency of spontaneous activity and the coordination between regions of the sensorimotor cortex (Figure 8). The experience-independent emergence and development of spontaneous S1-M1 co-activity at these young ages does align with the fact that this form of activity is not driven from the sensory periphery (Figure 7). This suggests that the initial establishment of this sensorimotor cortical network may be innate. Our experiments do not rule out though that the integration of this network into sensory-driven responses is experience-dependent at some point in development. Indeed, given that the activity-dependent maturation of each synaptic relay in sensory pathways occurs in sequence, with more peripheral connectivity developing earlier followed by thalamocortical and then local intra-cortical synapses, it may be that there is an experience-dependent phase later.

Overall, our findings suggest that there is an unexpectedly early establishment of functional connectivity between S1 and M1. There is preferential coordination of spontaneous activity in these regions from neonatal ages. At the same time, sensory stimulation can efficiently drive activity in S1 but is not yet able to engage this broader functional network even though it exists. Indeed, the spontaneous S1-M1 co-activity is largely not driven from the sensory periphery and develops independent of sensory experience, in contrast to the ascending pathways. This suggests that there is parallel, independent neonatal maturation of ascending sensory pathways and intra-cortical networks. We predict that the sensory-evoked activation of the full sensorimotor network emerges through maturation of a key, as yet unknown, synaptic pathway that a links the ascending pathway to the already-established S1-M1 connectivity.

## Methods

### Animals

All procedures were carried out in accordance with UK Home Office guidelines set out in the Animals (Scientific Procedures) Act 1986.

Mice expressing GCaMP6f in excitatory cortical neurons were generated by crossing two transgenic lines: Emx1-IRES-cre (005628 - The Jackson Laboratory, USA) and loxP-GCaMP6f-loxP (Ai95D) (028865 - The Jackson Laboratory, USA). Experiments were performed on pups of both sexes from ages P1 to P9. In early stages of the project, using simple observation of new-born pups under fluorescence illumination, we noted seemingly off-target expression of the indicator across the entire body in a subset of animals. In contrast, other animals had expression that was restricted to the brain. We compared the distribution of expression in the brains of these two types of mice. Histological investigation of ‘whole body expression’ pups showed widespread GCaMP6 expression, including in sub-cortical regions of the brain (Supplementary Figure 1a – body). In contrast, in the “brain-only” animals, GCaMP6 expression was restricted to the neocortex as expected (Supplementary Figure 1a – brain only). Retrospective comparison of parental genotype linked “whole body expression” in some pups to double-transgenic sires. This suggested there may be germline recombination within some sperm of males carrying both Emx1-IRES-cre and Ai95D transgenes (Luo et al., 2020). In line with this, a breeding strategy using only singly transgenic males yielded no off-target “whole body” expression in pups. This strategy was adopted for all subsequent experiments.

### Surgical procedure and anaesthesia

Surgical anaesthesia was induced with inhalation isoflurane (2-3%) in medical oxygen. Body temperature was maintained at 37^0^C during surgery and imaging, using a heatmap (Harvard Apparatus, UK). Local anaesthesia (20µl 2% xylocaine with adrenaline – AstraZeneca, UK) was administered subcutaneously under the scalp. The scalp was removed and clear dental cement (C&B super bond kit - Prestige Dental UK, Bradford, UK) was applied to the exposed skull and to attach a head-fixation bolt (4-40 ¼” stainless steel set screw - Thorlabs Inc. USA) attached over the cerebellar plate. Animals were maintained at 37^0^C and breathing air throughout recovery from anaesthesia and imaging protocols.

In line with previous studies, we found cortical activity in neonatal mice was almost completely suppressed during the administration of isoflurane anaesthesia (Ackman et al., 2012; Adelsberger et al., 2005; Hanganu et al., 2006; Siegel et al., 2012). Spontaneous cortical activity started to appear a few minutes after cessation of isoflurane exposure (Supplementary Figure 1c-d). To further assess any lingering effects of anaesthesia, we measured the frequency of spontaneous events across the entire cortical field of view in 3 minute epochs at 30, 60 and 90 minutes after removal of isoflurane. The frequency of spontaneous activity significantly increased between 30 minutes and 60 minutes post-anaesthesia, but there was no difference between 60 minutes and 90 minutes (Supplementary Figure 1c). These results suggest that although absolute suppression of spontaneous cortical activity by anaesthesia subsides within minutes, there may be a longer-term effect that takes up to an hour to resolve in younger animals. As such, all imaging data analysed was collected at least 60 minutes after removal of isoflurane.

### Widefield calcium imaging protocol

Awake animals were attached to an articulated ball and socket head mount. Imaging was performed on a tandem lens (50mm, 1.4f - Sigma Imaging, UK) fluorescence macroscope, with a 470 nm blue LED excitation light (M470L3 – Thorlabs Inc. USA) and 500nm long pass emissions filter. Images were captured with CMOS camera (Q Imaging, Canada) as 12-bit 960×540 pixel TIFF files at 50Hz frame rate. Movement was recorded using a piezo bender (Piezo Systems Inc, USA) placed under the animals’ body. Deviations in measurements from the pressure sensor correlated well with motion detected in video recordings of the limbs confirming they are a good indicator of gross body movements (Supplementary Figure 2ai). To isolate periods of movement, piezo voltage recordings were binarised using a thresholded envelope to define periods of movement and rest (Supplementary Figure 2a).

Images were collected in 10,000 frame epochs (3.3 minutes). Multiple stimulated and spontaneous recordings were collected for each animal, with a 30-minute interval between epochs of each type.

For single whisker stimulation individual whiskers were sequentially threaded into a glass capillary tube attached to a piezo bender (Q220-A4-103YB - Piezo Systems Inc, USA) to 1mm from the snout and a single deflection of 100ms and 150 µm displacement was delivered every 10 seconds, with 20 repeats. For whisker array stimulation a custom 15×8mm plastic paddle attached to a 9V servo motor (SG92R – Tower Pro) was used to displace the whole whisker field in a caudal to rostral direction for a 30ms displacement, at a 10 second interstimulus interval, for 20 repeats.

### Sensory manipulations

For whisker trimming experiments all whiskers on the left side were removed daily from day of birth until experiments were carried out at P7. The procedure was performed on awake, scruffed animals using microdissection scissors (World Precision Instruments, USA) to cut the whisker down to the snout. For acute peripheral silencing 30µl of 2% xylocaine (AstraZeneca, UK) was injected subcutaneously into the right whisker pad.

### Histology

Brains were removed from the skull and drop-fixed in 4% paraformaldehyde made in Dulbecco’s PBS (Sigma-Aldrich Ltd, UK) for 48hrs at 4^0^C and then transferred to Dulbecco’s PBS for storage. Brains were sectioned at 50µm from frontal to occipital pole using a vibratome (Leica VT1200 - Leica Microsystems, UK). Sections were mounted with Vectashield with DAPI medium (Vector Laboratories Ltd, UK) and visualised with a fluorescence microscope (DM IRB - Leica Microsystems, UK) and captured as tile scanned 696×52- 8 bit .lif images at 5x magnification.

### Image analysis

Imagine data were analysed using custom designed software in MATLAB (Mathworks, MA, USA). Images were imported and underwent bilinear transformation to reduce the spatial resolution to 480×270 pixels.

Periods of movement were detected from the 1kHz piezo bender timeseries (Supplementary figure 3). A 20x averaging temporal filter was applied to match the frequency of calcium image capture. Matlab’s ‘envelope’ function was used to delineate a continuous, positive time course of movement. A threshold was assigned manual for each trace and periods that exceeded this were assigned as movement. This binary movement log was used in imaging analysis to investigate periods of quiet rest.

Cortical regions of interest (ROIs) were manually delineated each animal. The area of activation evoked by whisker stimulation was used as a known anchor point for each animal (Figure 2d) and then both the Allen Institute’s adult mouse atlas and the developing cortex atlas defined by (Ackman et al., 2014) were used to assign the location of other cortical regions. For spontaneous recordings average fluorescence timeseries for these regions were calculated and baseline corrected using the lowest 5% of values to produce Δf/f traces, that were then filtered with a 7Hz lowpass Butterworth filter to remove heartbeat-associated fluctuations (Supplementary Figure 1d). For spontaneous activity frequency events were detected using an automated peak detection algorithm with a threshold of 1% Δf/f (twice the value of average noise) from local baseline, with events occurring during periods of movement discarded (Figure 2b&e). Correlations were calculated using Pearson’s correlation between corrected timeseries (Figure 2c & f).

Seed pixel maps (SPMs) were generated using pairwise Pearson’s correlation between baseline and temporally corrected timeseries of a single pixel in the barrel cortex and every other pixel in the cortex (Figure 3a). S1 and M1 areas of activation in these SPM were outlined and the average timeseries calculated. These were used, along with average visual cortex (V1) timeseries, to calculate the coincidence ratio. Events in all timeseries were detected with the automatic peak algorithm used previously. At each timepoint of an event in S1 occurring during rest a sliding lag window was applied to the other regions to determine if an event simultaneously had occurred, and the ratio of occurrence calculated (Figure 3c). For all events detected in S1 the image frames at the peak were manually assessed and categories by whether there was a change in Δf/f in M1 accompanying S1 activity (Figure 4b).

Spatial activation was calculated by averaging the images of 1 second following deflection for all stimulations in which no movement was detected in this period. The average heatmap from an epoch of recording was used to calculate the ROI of activation was the area around the peak Δf/f that was 50% of the maximum (Figures 5a & 6a). This ROI was used to extract an average timeseries and was baseline corrected to the 500ms of rest preceding each deflection. Amplitudes of events were calculated as the difference between the peak and the minimum of the preceding 500ms (Figure 6d).

### Non-negative matrix factorization (NMF)

Image stacks were geometrically aligned to a custom 2D projection of the Allen Common Coordinate Framework v3 (CCF), within and across recordings, using nine anatomical landmarks: the left, centre, and right points where anterior cortex meets the olfactory bulbs; the sagittal sinus midline; bregma; and the centre of mass of the somatosensory and visual cortex from both hemispheres. After alignment and registration, non-neural pixels were masked using the outline of the atlas 2D projection, and recordings were spatially binned to 68×68 pixels (104 μm^2^/pixel) and the ΔF/F over time was computed individually per pixel. Since the factorization requires non-negative pixel values, the recording was normalized to a range of 0 to 1 using the maximum and minimum pixel values per recording. The data comprises 4 different developmental ages (P3, P5, P7 and P9), 6-7 animals per age, and 2 recordings per animal.

We used a standard non-negative matrix factorization (NMF) algorithm to discover spatial motifs in widefield data. The minimally pre-processed data, *X* (*P* pixels by *T* time points), is factorized into the 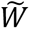 (spatial motif) and 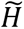 (temporal weights of the motif) factors which minimize the following cost function to produce an optimal reconstruction 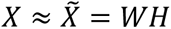:

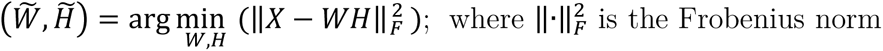

This problem is optimized using gradient descent with multiplicative updates (Mackevicius et al. 2019, eLife). Each optimization is run multiple rounds to assess the stability/consistency of each model. Ten independent NMF fits were run with different initial conditions for each motif number (from 1-30) and we choose the case that yields a factorization that explains on average ≥ 75% of the variance (EV) from the original dataset:

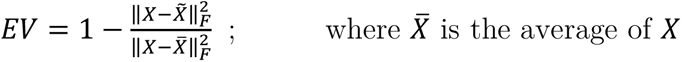

The motifs obtained per NMF fit from the same recording were hierarchically clustered to obtain the set of spatial motifs per recording (Supplementary figure 2). For the clustering, motifs were renormalized to 0 to 1 and spatiotemporally smoothed with a 2D Gaussian filter (*σ* = [1, 1]). The 2-D correlation coefficient between each pair of spatial motifs from the same recording was computed, and the distances between each pair of observations was used to create a hierarchical cluster tree. The maximum number of clusters was set to the number of NMF motifs originally found at 75% EV. Once the set of spatial motifs per recording was generated, the temporal weights of each motif was recalculated, fixing *W* to these set of spatial motifs.

To investigate whether these motifs could reflect underlying computations, we correlated (Pearson correlation) the temporal weightings of each motif in the set with the corresponding movement (piezo sensor) trace.

The presence of the S1-M1 co-activity motif within the set of spatial motifs per recording was visually assessed. Each data point represents the observation of a S1-M1 co-activity motif, with the possibility of occurring more than once in the same recording. The contribution of the S1-M1 co-activity motif per recording was calculated as the average of the Relative Temporal Contribution (RTC) of the motif across the recording:

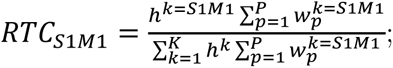

where *K* is the total number of motifs per recording, *P* is the total number of pixels, *w^k^* and *h^k^* are the spatial and temporal weight vectors for the *k^th^* motif.

### Statistical analysis

Statistical analysis was performed using R version 3.5.0 (The R Project). Data were checked for normality of distribution using both a Shapiro-Wilk’s test and ANOVA or T-testing, or their non-parametric alternatives, were used where appropriate. Boxplots show median (line), interquartile range (box) and data extremes (whiskers).

## Author Contributions

CMC designed and performed experiments, analysed data and contributed to the manuscript writing. LMS designed and implemented data analysis and contributed to writing the manuscript. NW supervised data analysis and contributed to the manuscript. KL supervised CMC and contributed to project design. MCA designed and supervised the project, built imaging hardware, performed data analysis and contributed to manuscript writing.

## Acknowledgments

Funding for the project was provided by Medical Research Council (1514380), Wellcome Trust (220102/Z/20/Z) and EUFP17 Marie Curie Actions (PCIG10-GA-2011-303680). NW is supported by a Turing Fellowship from the Alan Turing Institute.

