## Supplemental Figure 1 for "Early functional connectivity in the developing sensorimotor network that is independent of sensory experience"

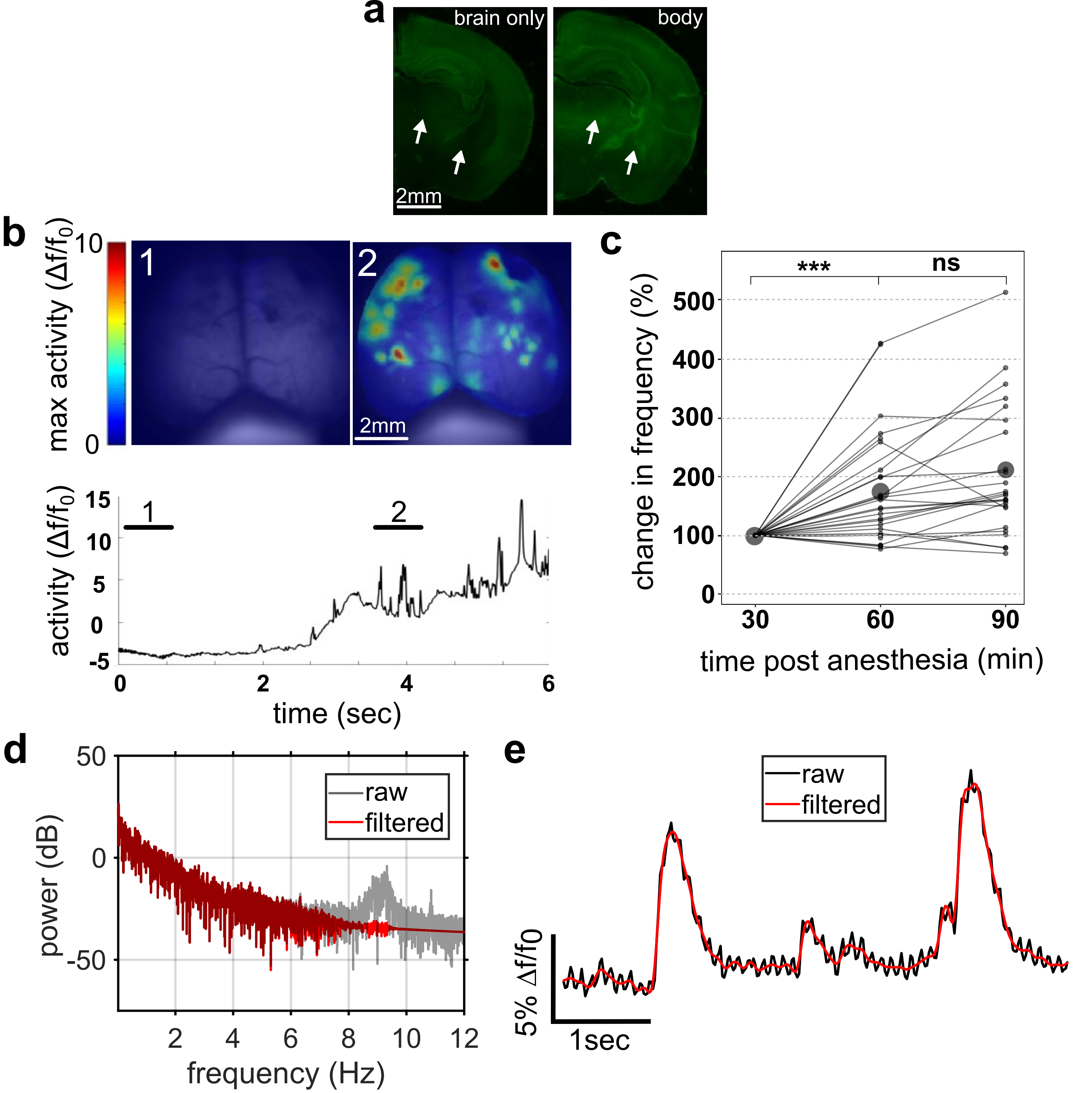


**Supplementary Figure 1. In vivo widefield calcium imaging in postnatally developing mice.**

a. Coronal brain sections of P7 animals with on-target cortical (brain-only) and off-target (body) expression. Note the different expression patterns in subcortical areas labelled with arrows.

b. Time-course of recovery of spontaneous cortical activity following removal of surgical (isoflurane) anaesthesia in a P7 mouse. Maximum activity projection maps for periods indicated on trace of fluroescence intensity changes across cortex.

c. Frequency of spontaneous cortical activity events at 30-min intervals post-anaesthesia, normalised to first session. Activity significant increases between 30 and 60 seconds after anesthesia cessation ($p < 0.001$, two-way repeated measures ANOVA).

d. Power spectrum of raw spontaneous fluorescence intensity changes across 3 minute recording in barrel cortex of P5 mouse showing characteristic peak in power at ~8-10Hz that corresponds with the heartbeat of neonatal rodents (black trace). Lowpass filtering below 7Hz removes this activity while preserving lower frequency activity (red trace).

e. Segment of spontaneous fluorescence changes (from recording used in d) before (black trace) and after (red trace) low-pass filtering, which removes the high frequency oscillation leaving behind the slower fluorescence transients.
