## Supplemental Figure 2 for "Early functional connectivity in the developing sensorimotor network that is independent of sensory experience"

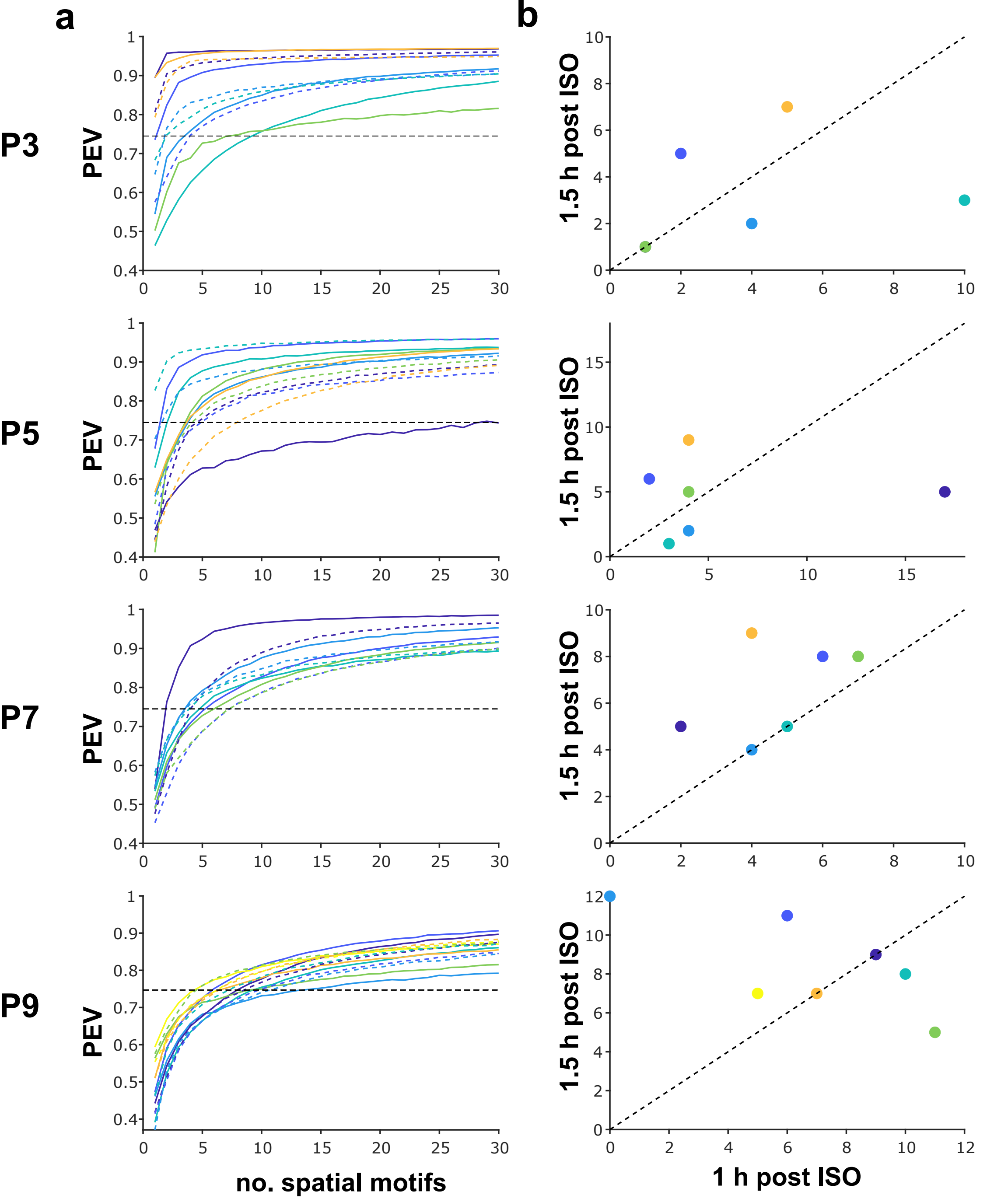


**Supplementary Figure 2 - NMF output for each recording.**

a. Precentage Explained Variance (PEV) of individual recordings by varying numbers of motifs specified by NMF at each age (P3, P5, P7, P9). Dashed line shows the 75% PEV level used to compare motif contributions.

b. Comparison of the number of moitfs required to reach 75% PEV in the two recordings from each animal. First recording was 1hr post-isoflurane, with second recording 1.5hr post isoflurane. There is variability in number of motifs required to explain variance across sessions in each animal but no systematic effect of post-anaesthesia duration.
