## Supplemental Figure 3 for "Early functional connectivity in the developing sensorimotor network that is independent of sensory experience"

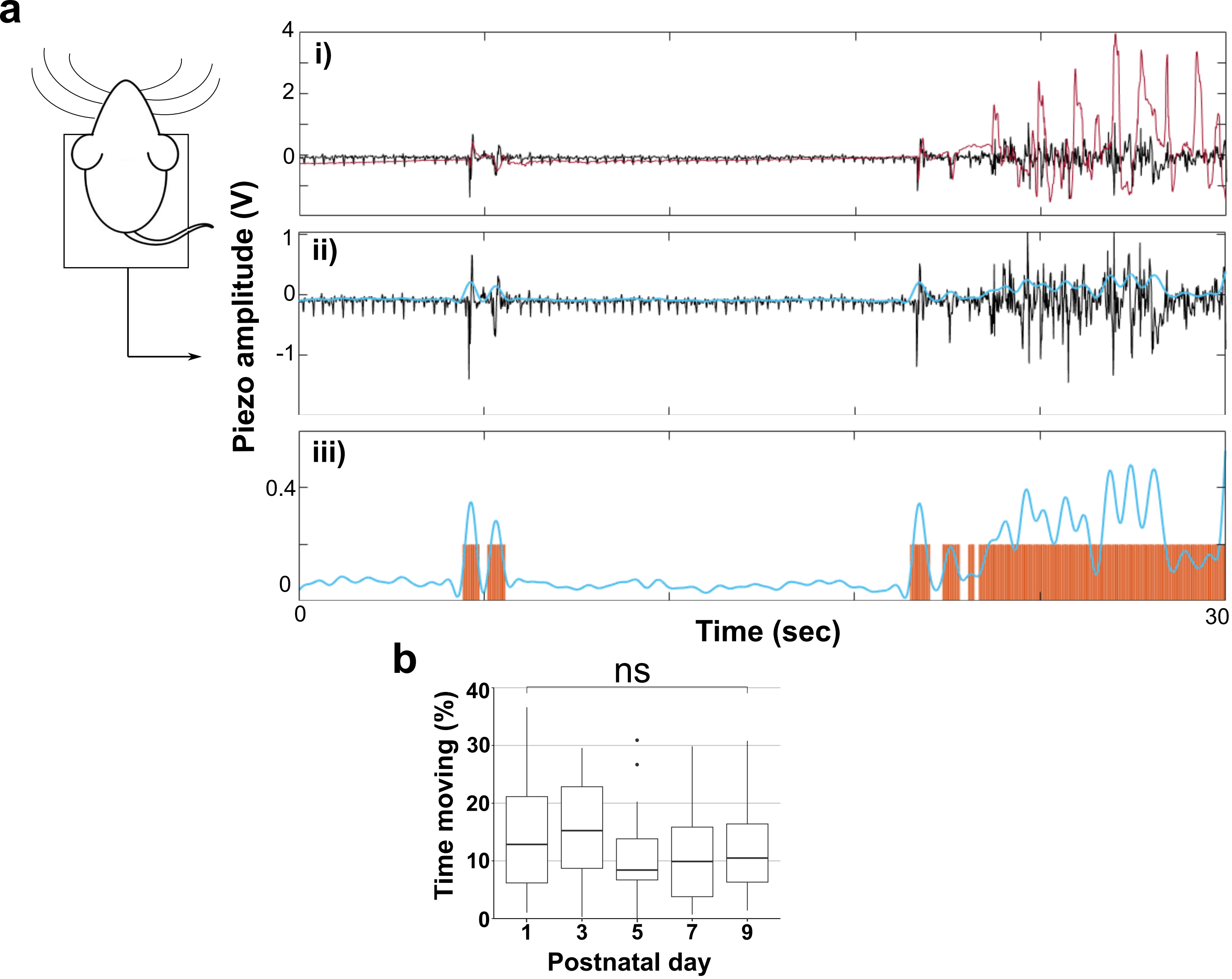


**Supplementary figure 3. Body movements could be reliably detected using a pressure sensor under the body.**

a. Movement was recorded from a piezo bender under the mouse's body. i) In preliminary tests these movement traces (black) were compared with video recordings of the body, showing reliable capture of movement as deflections in the piezo output voltage. ii) An automated envelope (blue) was created around to give a smoothed output of movement. iii) From this envelope (blue) a binary log (orange) of movement vs rest could be calculated.

b. Percentage time spending moving during spontaneous recordings did not change with developmental age.
